# Mechanistic and thermodynamic characterization of antivirals targeting druggable pocket of SARS-CoV-2 nucleocapsid

**DOI:** 10.1101/2022.03.12.484092

**Authors:** Preeti Dhaka, Ankur Singh, Shweta Choudhary, Rama Krishna Peddinti, Pravindra Kumar, Gaurav Kumar Sharma, Shailly Tomar

**Affiliations:** Department of Biosciences and Bioengineering, Indian Institute of Technology Roorkee, Roorkee, Uttarakhand - 247667, India; Department of Chemistry, Indian Institute of Technology Roorkee, Roorkee, Uttarakhand-247667, India; Indian Veterinary Research Institute, Izatnagar, Bareilly, Uttar Pradesh state - 243122, India

**Keywords:** SARS-CoV-2, Nucleocapsid, RNA binding domain, Drug repurposing, NTD

## Abstract

The N-terminal (NTD) and the C-terminal (CTD) domains comprises the structure of the SARS-CoV-2 Nucleocapsid (N) protein. Crystal structure of the SARS-CoV-2 N protein determined by Kang et al, 2020, reveals the N-terminal RNA binding domain as a unique drug binding site. The present study targets this unique pocket with identified antivirals using structure-based drug repurposing approach. The high-affinity binding of potential molecules was characterised thermodynamically using Isothermal titration calorimetry. The selected molecules showed an inhibitory RNA binding potential between 8.8 μM and 15.7 μM IC_50_ when evaluated with a fluorescent-based assay. Furthermore, in an in vitro cell-based antiviral assay, these ten antiviral molecules demonstrated high effectiveness in halting SARS-CoV-2 replication. Telmisartan and BMS-189453, the two highly potent antivirals, have ∼0.98μM and 1.02 μM EC_50_ values with the selective index of >102, and >98, respectively. For the first time, this study presents drug molecules specifically targeting the NTD of SARS-CoV-2, offering essential insights for the development of therapeutic interventions against this virus, which is still a potential global threat to public health.

## 1. Introduction

Since the outbreak of the severe acute respiratory syndrome coronavirus 2 (SARS-CoV-2) pandemic (COVID-19), efforts have been made to create coronavirus countermeasures. The mRNA vaccines, adenovirus vectors, inactivated viruses, subunit vaccines are now available, and number of antiviral drugs were used during pandemic to reduce mortality and hospitalization. However, still there is a need for novel effective antivirals to combat emerging variants of SARS-CoV-2 (Omer et al., 2023). SARS-CoV-2 infection in humans can cause severe symptoms such as pneumonia, acute respiratory distress syndrome, cytokine storm and organ failure, highlighting the need for effective antiviral therapy (Masters, 2006). The viral genome consists of four structural proteins including the Nucleocapsid (N) protein essential for regulating viral RNA replication, host cell metabolism, and viral assembly (Hurst et al., 2009; V et al., 2003). The N-terminal RNA encapsidation domain (NTD; residues 46-174) and the C-terminal oligomerization domain (CTD; residues 247-364) are two functional domains of N-protein, connected by a central linker region. Research on the SARS-CoV-2 N-protein indicates that the NTD is highly conserved and has structural and functional similarities with other coronaviruses (V’kovski et al., 2021). Viral RNA and positively charged NTD form a ribonucleoprotein (RNP) complex through hydrophobic and electrostatic interactions (CK et al., 2009). The crystal structures of NTD-N protein of coronaviruses reveals the RNA binding pocket that is primarily responsible for encapsidation of viral RNA. Targeting a specific RNA-binding pocket on the N-protein could potentially disrupt the interaction and serve as a promising target for the development of anti-COVID therapeutics (Kang et al., 2020). The process of establishing a new medication in the pharmaceutical market involves costly and time-consuming pre-clinical and clinical trial stages. Efficient screening of existing compound libraries against current and upcoming diseases is crucial to expedite drug discovery and development (Meinhardt et al., 2020; Montes-Grajales et al., 2020; Rani et al., 2022). The process can be sped up by using various drug libraries for structure-based screening, molecular dynamic (MD) simulation, biophysical and biochemical validation followed by *in vitro* testing of molecules against target proteins (Mudgal et al., 2020; Rani et al., 2022). Additionally, the development of a high throughput fluorescence-based protein-RNA binding assay helps in evaluating impact of investigational drugs on molecular interactions and provides a inhibition kinetics method (Byun et al., 2020; Hwang et al., 2011). This study aimed to identify small molecules from three different libraries that could potentially target the NTD-N protein and inhibit the replication of SARS-CoV-2. Twelve promising molecules were identified through high-throughput drug screening and their binding affinities and inhibitory effects on NTD-RNA interactions were evaluated using isothermal titration calorimetry (ITC) and fluorescence intensity-based assays respectively. Ten molecules were identified as inhibitors of SARS-CoV-2 in cell-based investigations. The study provides a strong foundation towards development of antiviral treatment to combat emerging SARS-CoV-2 and its variants.

## 2. Materials and Methods

### 2.1. Production of SARS-CoV-2 NTD protein

Gene fragment encoding the NTD was PCR amplified using the template plasmid received from National Centre for Cell Science (NCCS), Pune (a kind gift from Dr Janesh Kumar). The gene was cloned in pET28c vector using 5’-ATATGGCTAGCCGGCCCCAAGGTTTACCCAATAATACTGC-3’ as forward primer having *Nhe*I restriction site & 5’-AAGCTTGTCGACTTATTCTGCGTAGAAGCCTTTTGGCAA-3’ as reverse primer having *Sal*I restriction site. The cloned plasmid was transformed in *E. coli* DH5_α_cells, followed by plasmid isolation, and gene insert confirmation through DNA sequencing. Expression and purification of protein was then performed using Rosetta (DE3) *E. coli* as previously described (Kang et al., 2020). The molecular weight ∼21.7 kDa and purity of the 6xHis-tagged NTD protein was confirmed on sodium dodecyl sulphate–polyacrylamide gel electrophoresis (SDS-PAGE). The purified protein was dialysed using 1XPBS buffer (pH 7.3) at 4 °C and concentrated to ∼3.6 mg/ml.

### 2.2. Virtual screening, molecular docking, and MD simulation studies

The structural superimposition of SARS-CoV-2 NTD (Protein Data Bank, PDB ID: 6M3M) (Kang et al., 2020) with NTD of SARS-CoV (PDB ID: 2OFZ) (Saikatendu et al., 2007), MERS-CoV (PDB ID: 4UD1) (Papageorgiou et al., 2016), and HCoV-OC43 (PDB ID: 4KJX) (Lin et al., 2014) were performed using PyMol (Delano, n.d.) to observe the conservation of residues involved in RNA binding (Figure S1). The Multiple sequence alignment (MSA) was done using MultAlin (Corpet, 1988) and visualized using ESPript3.0 (Robert and Gouet, 2014). For molecular docking study, the atomic structure of NTD (PDB ID: 6M3M) was retrieved from RCSB-PDB and 3D-structure of guanosine monophosphate (GMP) was extracted from the PubChem database. Docking studies were conducted targeting the grid box around RNA binding residues with dimensions (44Å×46 Å×43Å) and centre point coordinates (X= 13.5, Y= −6.2, and Z= −18). As a reference parameter, docking was performed in the same pocket for determining the binding energy (B.E) of GMP (−5.6 kcal/mol). In a similar manner, molecular docking was performed using PJ34, an inhibitor targeting the GMP binding pocket of NTD of HCoV-OC43 and reported to significantly inhibit the viral replication(Chang et al., 2016; Wu et al., 2023). The molecular docking of the best-fit conformation of PJ34 revealed the B.E. as −6.2 kcal/mol. Structure-based virtual screening of compound libraries including LOPAC^1280^ (“LOPAC®1280,” n.d.), natural compounds (“Natural Products,” 1998), and FDA-approved drugs (Liu et al., 2022) was performed using PyRx 0.8 (Dallakyan and Olson, 2015) and AutoDock tools/Vina (Morris et al., 2009; Trott and Olson, 2009) against RNA binding pocket of NTD protein. Molecules were energy minimized and converted to Autodock molecules (.pdbqt) format using PyRx 0.8 with OpenBabel. Top twelve molecules displaying B.E. higher than the NTD-GMP and NTD-PJ34 complex were further selected for detailed interaction analysis using PyMOL and LigPlot^+^ (Laskowski and Swindells, 2011).

MD simulation of NTD-inhibitor complexes was performed using GROMACS (Van Der Spoel et al., 2005) 2022.2 program for 100 ns on Linux based workstation. The program pdb2gmx was used to process the interacting poses of structures and the small molecules topology was generated using CHARMM36 (Huang et al., 2016) General Force Field (CGenFF) (Vanommeslaeghe et al., 2010). The method for MD simulation studies was followed as described previously (Rani et al., 2020). The data were processed using OriginLab 2018.5 (www.originlab.com).

### 2.3. Isothermal Titration Calorimetry (ITC)

ITC experiments were performed using MicroCal ITC_200_ microcalorimeter (Malvern, Northampton, MA) (“MicroCal iTC200 isothermal titration calorimeter from Malvern | Product support | Malvern Panalytical,” n.d.) in 1XPBS buffer at 25 °C with reference power 9 μcal/s. Protein and ligands were dissolved in 1XPBS buffer. For NTD-ligand binding experiments, the 100-400 μM concentration of ligands (in syringe) were titrated against 10-20 μM NTD protein (in cell). After equilibration, a total 20 injections of syringe samples were titrated in the cell with one injection of 0.4 μl followed by 19 subsequent injections of 2 μl, each spaced by 220 seconds. Using Malvern’s Origin 7.0 Microcal-ITC200 analysis programme, the one-site binding model evaluated the binding parameters to determine the reaction stoichiometry (n), enthalpy (H), entropy (S), and binding constants (Ka) values.

### 2.4. Fluorescence intensity-based NTD-RNA binding assay

The fluorescence intensity-based assay was performed to characterize the binding of RNA with NTD and to assess the inhibitory potential of identified molecules. Fluorescently labelled RNA (5’s 6-FAM-labeled (UCUCUAAACG) 10-mer RNA) was procured from Biotech Desk Pvt Ltd, India (“Welcome to Biotech Desk Pvt. Ltd.,” n.d.), and prepared its aliquot using DEPC treated nuclease free water. The binding buffer (20 mM HEPES, pH-7.0, and 100 mM NaCl) was prepared in DEPC-treated water. The purified NTD with varied concentration of 0 μM −10 μM was added to 96-well black flat bottom polystyrene microplates (Corning #3881) followed by addition of 1 nM RNA to each well and the plates were incubated at 4 °C for 30 min. Wells containing only RNA were treated as blank controls whereas the wells having non-binding protein (Main protease (Mpro) of SARS CoV-2) with RNA were treated as negative control. The fluorescence intensity was measured on a Synergy HTX multimode plate reader (Agilent BioTek) using Gen5 software at an excitation wavelength of 485/20nm and emission wavelength of 528/20nm with read height 8.5mm. Using the developed protocol, the inhibitory potential of molecules were assessed after incubating them with NTD followed by the addition of RNA. The RNA with NTD and no inhibitor were considered as positive controls. Fluorescence intensities were measured using plate reader and percentage inhibition vs concentration graphs for binding of NTD to RNA were plotted using Graph Pad Prism(“Prism - GraphPad,” n.d.). The data represents the average from a duplicate set of reactions.

### 2.5. Cell culture and virus preparation

Eagle’s Minimum Essential Medium (EMEM; MP Bio) was used for growing vero cells and supplemented it with 10% FBS (Himedia; India), 1 μg/ml Gentamycin (Himedia). Cells were maintained at 37 °C along with 5% CO_2_ in a humid environment. SARS-CoV-2/Human/IND/CAD1339/2020 was propagated and titrated using Vero cells. The virus isolate was genetically characterized by whole genome sequencing (GenBank accession no: MZ203529). The titer of the virus was evaluated using a 50% tissue culture infectious dose (TCID_50_/ml) assay (Reed and Muench, 1938), and virus stocks were kept at −80 °C. All studies on infectious SARS-CoV-2 were performed in Biosafety level 3 facility at Indian Veterinary Research Institute, Bareilly, after obtaining necessary approvals from Biosafety Committees.

### 2.6. Assessment of the antiviral efficacy of small molecules against NTD

Standard MTT assay was used to assess the cytotoxic effects of molecules on Vero cells as reported previously (Mudgal et al., 2020). Vero cells were first pre-treated for 2-hour with the variable compound concentrations followed by infection with SARS-CoV-2 at 0.01 multiplicity of infection (MOI) for 2-hour. The plate was washed and cells were incubated in fresh medium containing inhibitors. The plate was incubated for 48-hour followed by a single freezing–thaw cycle. Viral RNA extraction was done from lysates using the TRU-PCR viral RNA extraction kit (BlackBio) to the manufacturer’s specifications. With the objective of quantifying the virus yield as previously mentioned (Wang et al., 2020), quantitative real-time RT-PCR (qRT-PCR) was performed using the commercial COVISure-COVID-19 Real-Time PCR kit (Genetix) in compliance with the manufacturer’s instructions. Hydroxychloroquine, a reference drug, was utilised as a positive control. Half-maximum effective concentration (EC_50_) values were determined using GraphPad Prism’s non-linear regression fit model, and data was represented as the mean for duplicate sets of reactions. The TCID_50_ assay was used to confirm the antiviral potential of small compounds (Reed and Muench, 1938). Briefly, Vero cells were cultured overnight in a 96-well plate and infected with 10-folds consecutive dilutions of lysates from the antiviral assay. The presence and absence of the cytopathic effect (CPE) were observed during the 72-hour incubation period at 37 ºC and 5% CO_2_. After incubation, the cells were stained with 0.5% crystal violet and fixed with 10% paraformaldehyde for 6–8 hours. The dilution at which 50% of cells were infected was calculated and the data is reported as log_10_ TCID_50_/ml.

## 3. Results

### 3.1. Structure-based identification of NTD inhibitors

The crystal structures of NTD in complex with small molecules, including GMP from SARS-CoV and PJ34 inhibitor from HCoV-OC43, assisted in the identification of the druggable RNA binding pocket of coronaviruses (Lin et al., 2014; Papageorgiou et al., 2016; Saikatendu et al., 2007). Comparative sequence alignment and structural analysis showed that the RNA binding pocket in the NTD of SARS-CoV-2 N-protein is highly conserved among *Coronaviridae* family (Figure S1). Based on these sequence and structural comparisons, molecular docking of GMP targeting the RNA binding pocket was performed (Morris et al., 2009). Detailed molecular interaction analysis shows the participation of Ala56 and Tyr112 in H-bond formation with GMP, and Ala51, Ser52, Thr55, Ala91, Arg108, Tyr110, and Arg150 residues were observed to interact through hydrophobic interactions (Figure S2). The observed B.E. of the NTD-GMP complex was −5.9 kcal/mol. Furthermore, this B.E. was used as a guide to select potential small molecules from the selected compound libraries.

The screened molecules were docked using AutoDock Vina and AutoDockTools. The B.E. of these molecules were in the range −6.8 to −7.9 kcal/mol, which is significantly higher than the B.E. of NTD-GMP (−5.9 kcal/mol) (Fig. 1, Figure S2, and Figure S3). The binding conformations of the selected small molecules indicate that the residues involved in binding to GMP/RNA were also involved in interacting with these molecules (Fig. 1 and Figure S2). The comparative study of docked NTD-molecules complexes against NTD-GMP complexes reveals the involvement of some additional interacting residues of the same pocket that could possibly be responsible for increasing the binding affinity of the target protein for the small molecules (Table ST 1).

**Fig. 1.**
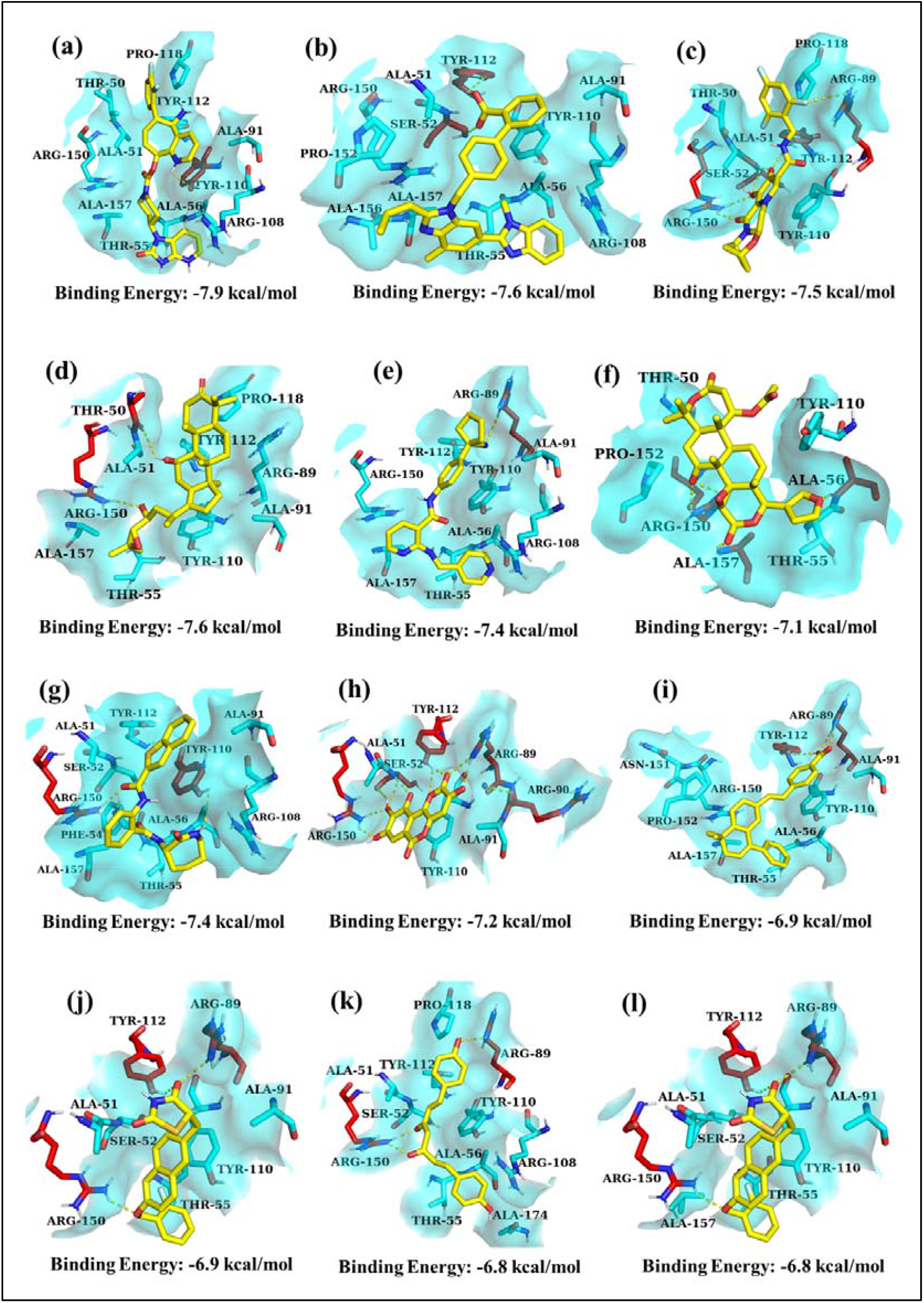
Molecular interactions of inhibitors in the RNA binding pocket of NTD of SARS-CoV-2 N protein using PyMol. (a) BMS-927711, (b) Telmisartan (MICARDIS), (c) Bictegravir, (d) Alisol_B, (e) Apatinib, (f) Nomilin,. (g) ANA-12, (h) Galloflavin_Potassium, (i) BMS-189453, (j) MCC-555, (k) Bisdemethoxycurcumin, and (l) Icilin occupying the binding pocket in a similar manner as GMP. Small molecules are shown in yellow coloured sticks, whereas the interacting residues of NTD are represented by cyan and red coloured sticks. The surface covered is indicated by cyan colour with 60% transparency. Yellow-coloured dotted lines show the intermolecular H-bond interactions.

FDA-approved molecules BMS-927711 makes one H-bond (Tyr110), Telmisartan makes two H-bonds (Ser52 and Tyr112), and Bictegravir shows six H-bonds, with four residues Ser52, Arg89, Tyr112, and Arg150. Natural product molecules Alisol_B interacts with two H-bonds (Thr50 and Arg150), Apatinib with one H-bond (Arg89), and Nomilin displays four H-bonds with residue Ala56, Arg150, and Ala157. LOPAC^1280^ molecules ANA-12 interacts with two H-bonds (Tyr110 and Arg150), Galloflavin_Potassium forms nine H-bonds with residues Ser52, Arg89, Arg90, Try112, and Arg150, BMS-189453 with two H-bonds (Arg89 and Tyr112), MCC-555, Icilin forms three H-bonds with residues Arg89, Tyr112, and Arg150, and Bisdemethoxycurcumin forms three H-bonds with residues Arg89 and Arg150 (Fig. 1, Figure S3, and ST 1). Further MD simulations were performed, and five parameters including the solvent accessible surface area (SASA), the radius of gyration (Rg), root mean square deviation (RMSD), root mean square fluctuation (RMSF), and intermolecular H-bonds for 100 ns were examined. The RMSD values indicated that majority of the complexes reach stability within 5-10 ns of simulation and falls within a range of 0.2 to 0.6 (Figure S4a). The RMSF values determine the fluctuation of the polypeptide chain as per C/span> atom coordinates from the reference. The increased RMSF values in residues 50-70 of the loop regions indicate enhanced flexibility, potentially facilitating small-molecules access to the RNA binding pocket (Figure S4b). According to the RMSF findings, NTD-ligand complexes have constant atomic mobility and as a result, ligands are expected to form stable complexes with NTD and, thus could prevent the binding of RNA to its pocket. Rg is employed to compute the distances between mass centres and the terminals of protein atoms. In this study, Rg plots showed that NTD-ligands complexes have comparable density to the NTD (Figure S4c). SASA is used to determine the solvation-free energy of every atom of the protein. The rise in the SASA plot suggests the conformational changes in the loop regions because of the presence of hydrophobic residues. SASA plots of complexes are equivalent to the NTD as shown in figure S4d. The number of H-bonds and their distribution increases with time, which represented that the system is attaining stability during simulation time, as shown in figure S4e. Furthermore, the binding of the selected small molecules has been validated by conducting ITC experiments using NTD protein.

### 3.2. Binding thermodynamic analysis of inhibitors targeting NTD

Using Origin 7.0 and Microcal-ITC200 analytical software, ITC isotherms were utilised and thermodynamic parameters were analysed using a one-site binding model for the non-linear curve. ITC analysis biophysically validated the good binding affinities of the identified molecules towards the purified NTD protein with *K*_*D*_ values in the range ∼5.0 to 90 μM (Fig. 2 and Table 1). Moreover, the N value is detected to be one for all the tested ligands that postulates their specificity for a single binding site of NTD (Table 1). The ΔH values obtained from ITC isotherms suggests that the binding of NTD with the selected ligands is exothermically driven (Table 1). In conclusion, the ITC titration curves clearly suggest that the identified molecules spontaneously binds to the NTD protein with suitable thermodynamic parameters and good affinities.

**Table 1.** Thermodynamic insights of top hit molecules binding to NTD Protein as obtained from ITC.

| Ligand | n | $K_D$<br>($\mu\text{M}$) | $K_A$ ($\text{M}^{-1}$) | $\Delta H$ (cal/mol) | $\Delta S$<br>(cal/mol/degree) |
| --- | --- | --- | --- | --- | --- |
| BMS-927711<br>(Rimegepant) | 1 | 13 | $(7.65 \pm 1.14)10^4$ | $(-17.70 \pm 1.87)10^4$ | -571 |
| Telmisartan<br>(MICARDIS) | 1 | 44.20 | $(2.26)10^4 \pm 561$ | $(-2.54)10^5 \pm 3787$ | -835 |
| Bictegravir | 1 | 22 | $(4.52)10^4 \pm (3.64)10^3$ | $(-6.56)10^4 \pm 2483$ | -199 |
| Alisol_B | 1 | 5.95 | $(1.68)10^5 \pm (3.83)10^4$ | $(-3.52)10^5 \pm (2.92)10^4$ | $(-1.16)10^3$ |
| Apatinib | 1 | 21.69 | $(4.61 \pm 1.20)10^4$ | $(-2.16)10^4 \pm 1977$ | -51.3 |
| Nomilin | 1 | 90 | $(1.11)10^4 \pm 312$ | $(-1.37)10^5 \pm 2796$ | -444 |
| ANA-12 | 1 | 40.40 | $(2.47)10^4 \pm 712$ | $(-1.81)10^5 \pm 3054$ | -590 |
| Galloflavin_Potassium | 1 | 36 | $(2.72)10^4 \pm (1.67)10^3$ | $(-2.01)10^5 \pm 6238$ | -654 |
| BMS-189453 | 1 | 39.21 | $(2.55)10^4 \pm 711$ | $(-4.09)10^5 \pm 6604$ | $(-1.35)10^3$ |
| MCC-555 | 1 | 27.4 | $(3.65)10^4 \pm (1.06)10^3$ | $(-3.02)10^5 \pm 4464$ | -993 |
| Bisdemethoxycurcumin | 1 | 82.6 | $(1.21)10^4 \pm 384$ | $(-1.63)10^5 \pm 3675$ | -529 |
| Icilin | 1 | 26 | $(3.83)10^4 \pm (5.70)10^3$ | $(-45.66 \pm 3.30)10^4$ | $(-1.51)10^3$ |

**Fig. 2.**
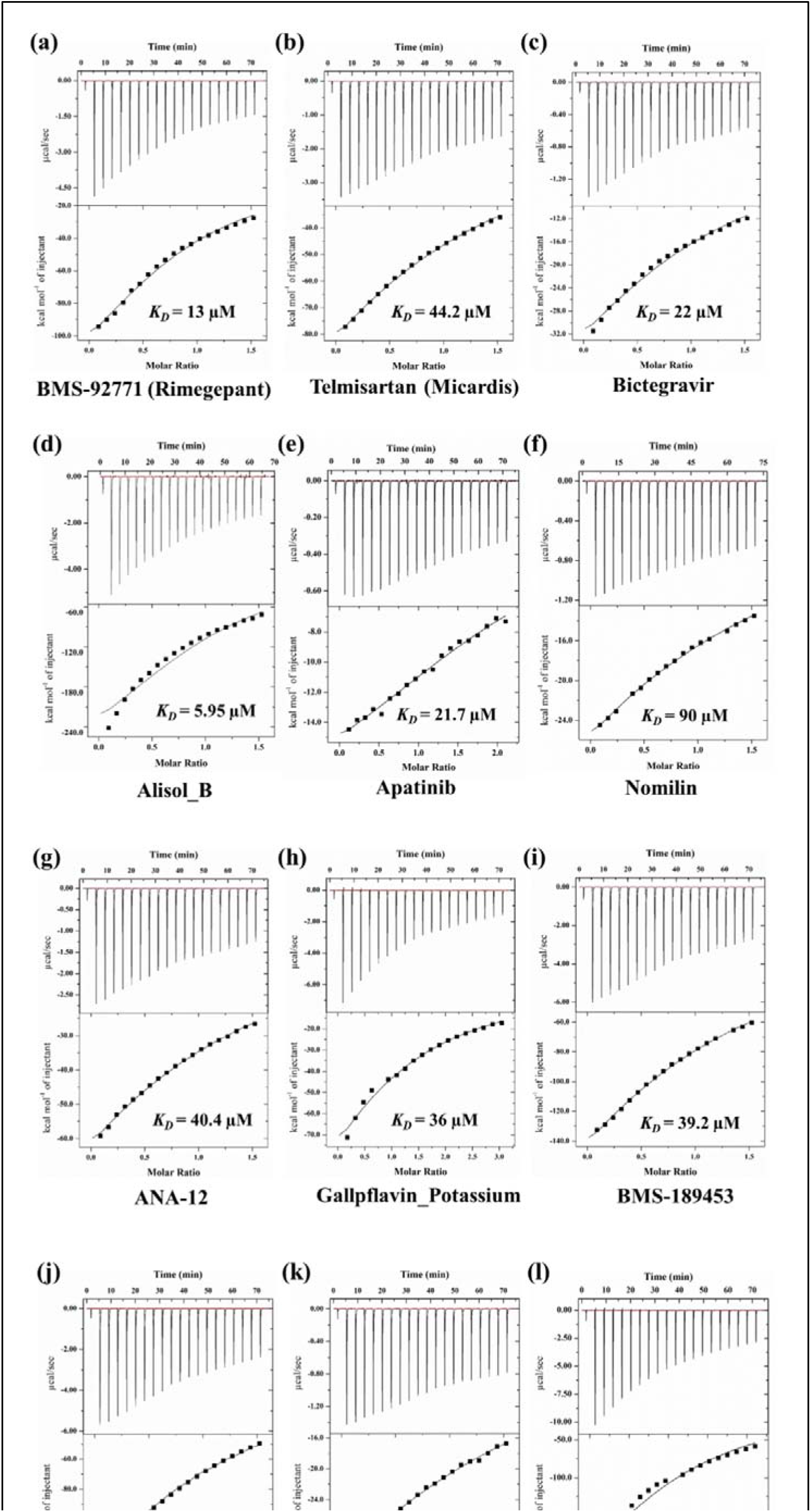
Binding isotherms of inhibitory molecules against the purified NTD protein using ITC

### 3.3. Fluorescence intensity-based NTD-RNA binding assay

Further, to determine the effectiveness of small molecules in inhibiting the targeted RNA-NTD molecular interactions, a fluorescence-based RNA binding inhibition assay was performed. The measured fluorescence intensity is directly proportional to the concentration of NTD protein, titrated with 1 nM of FAM-labelled RNA (Figure S5). Small molecules with the NTD-RNA binding inhibitory capacity lead to the decrease in fluorescence intensity value as compared to that of the free fluorescent RNA (Fig. 3). The half maximal inhibitory concentration (IC_50_) of twelve molecules were analysed using GraphPad Prism software. The identified molecules significantly inhibited the RNA binding activity of NTD with IC_50_ values in the range of 8 μM −16 μM (Fig. 3).

**Fig. 3.**
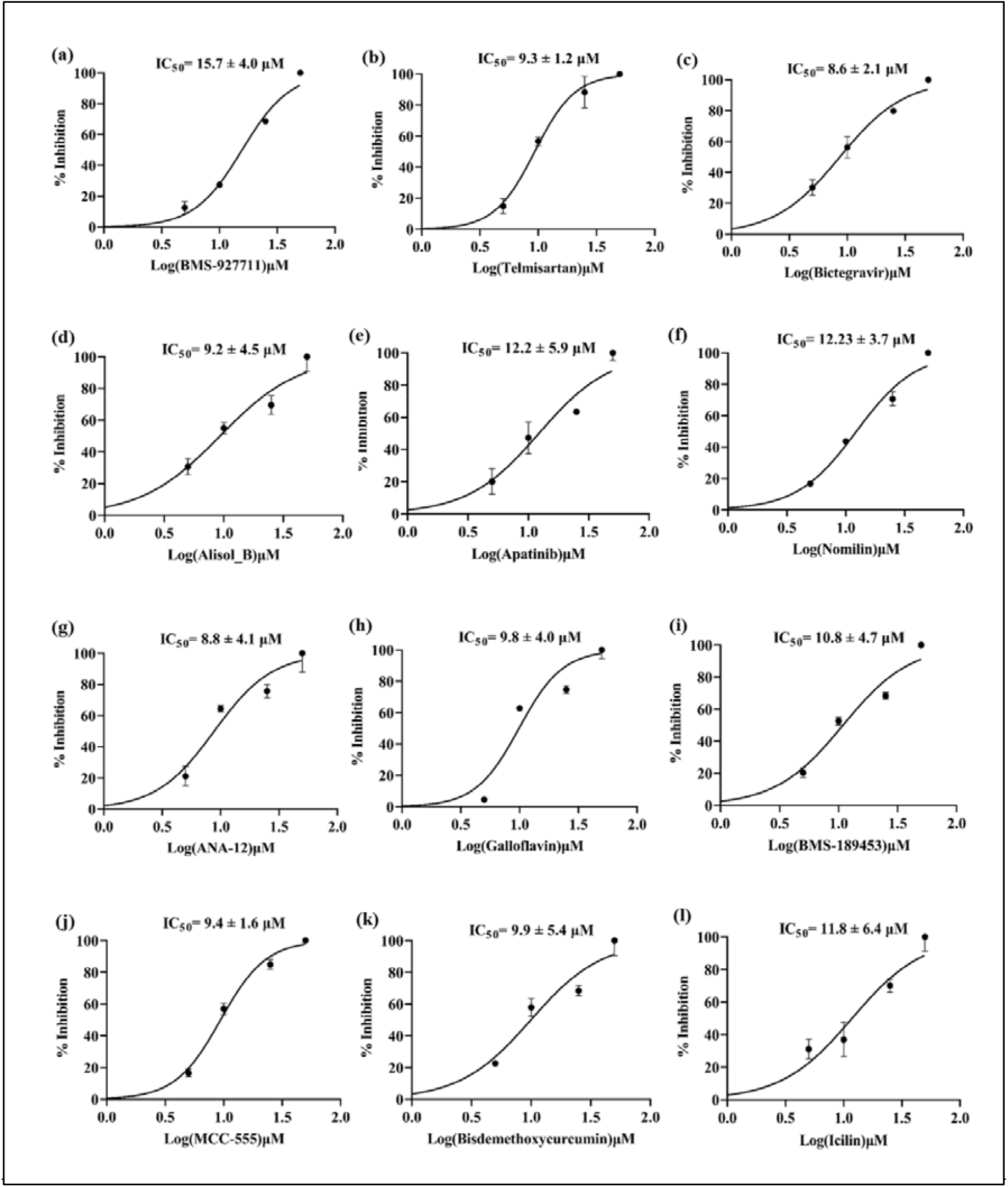
Fluorescence intensity-based NTD-RNA binding assay. The dose-response curves for percentage inhibition of NTD-RNA binding versus the concentrations of molecules plotted using GraphPad Prism software. The error bar of results represent the standard deviation from duplicate measurements.

### 3.4. Antiviral efficacy of small molecules against SARS-CoV-2

The antiviral efficacy of selected molecules was investigated using qRT-PCR and *in vitro* cell-based antiviral assays at non-toxic concentrations against SARS-CoV-2. As expected, excellent viral inhibition was observed for twelve molecules with a percentage inhibition of more than 90% at low concentrations and higher concentrations were able to completely abort the replication of virus. Telmisartan and BMS-189453 with EC_50_ ∼1.0 μM and ∼0.98 μM, respectively were found to be the best antivirals targeting NTD (Fig. 4). The results of qRT-PCR antiviral assays were validated by performing the TCID_50_ assay, where the compound-treated groups showed a significant decrease in viral titer than the untreated virus control. The TCID_50_/ml value observed for compound-treated groups was 10^2.49^ for MCC555 at 25 μM, 10^1.0^ for BMS-189453 at 12.5 μM, 10^1.8^ for Bictegravir at 25 μM, 10^2.03^ for Bisdemethoxycurcumin at 10 μM, and 10^2.16^ for ANA-12 at 20 μM in comparison to the virus control group (10^4.34^ TCID_50_/ml).

**Fig. 4.**
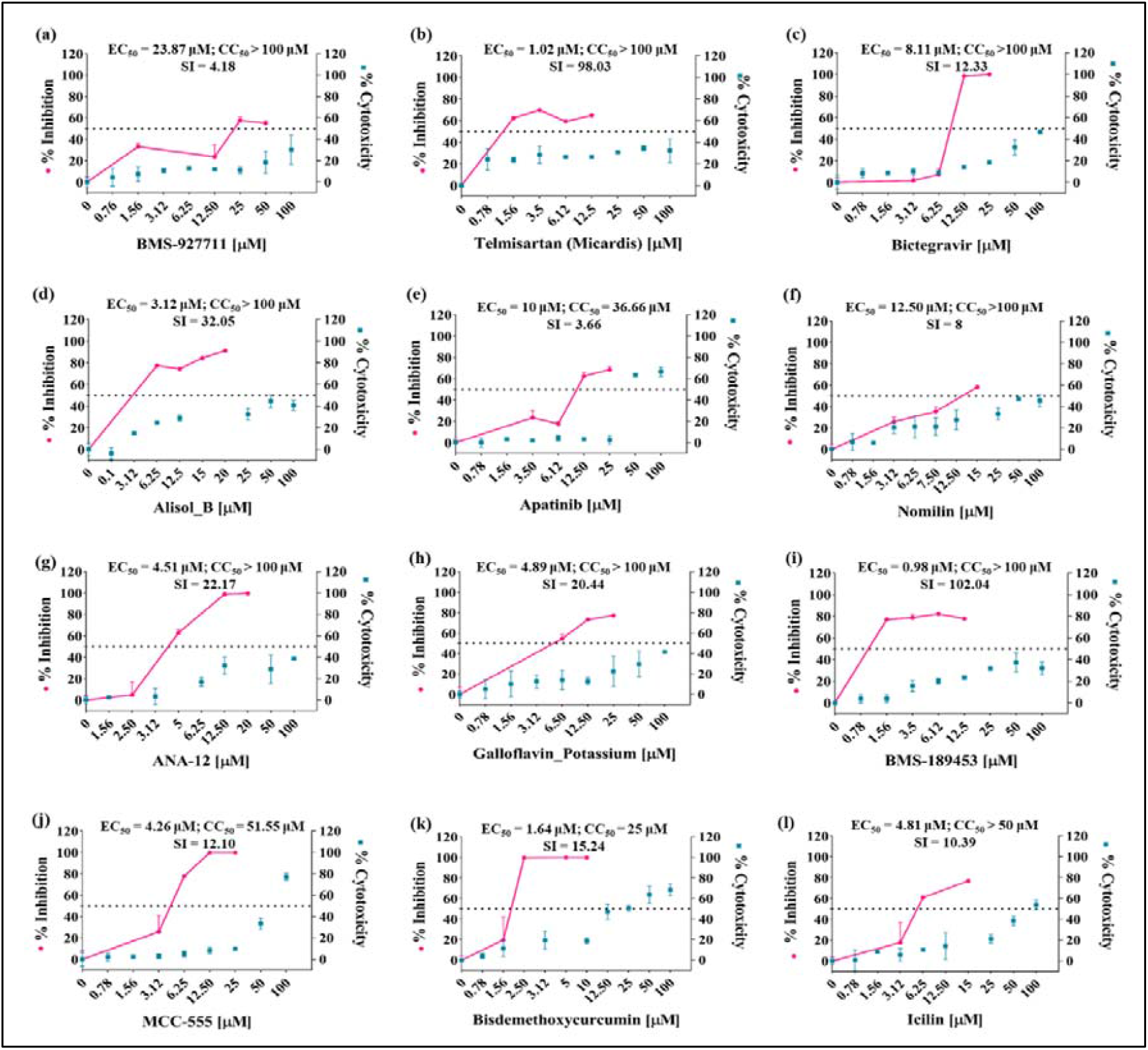
Anti-SARS-CoV-2 profiles of selected molecules. Percent cytotoxicity and percent inhibition graphs for identified molecules as determined by MTT assay and qRT-PCR experiments respectively. The pink circles specify the percentage inhibition of SARS-CoV-2 on treatment with selected molecules, and blue squares denote percentage cytotoxicity values. Results are representative of n = 2 and are shown as mean ± standard deviation. 50% cytotoxic concentration (CC_50)_, EC_50,_ and Selectivity Index (SI) values of each compound are displayed on upper panel of each corresponding graph.

## 4. Discussion

Identification of drug targeting sites in viral proteins is very crucial for the development of antiviral therapies. This study utilized the structural information available on the NTD of SARS-CoV-2 N-protein for identification of antivirals using the drug molecule repositioning approach. Biophysical binding characterization, biochemical RNA binding assays, and antiviral cell-based studies were performed to validate the inhibitory potential of identified molecules against the RNA-NTD molecular interactions. The N-protein plays a crucial role in viral assembly, genome packaging, replication, transcription, and host immune response regulation by forming a RNP complex (Bai et al., 2021). Previous studies suggest that small molecules targeting the RNA-binding pocket of the N-terminal domain could suppress viral infection by hindering the interaction between the N-protein and viral RNA (Lin et al., 2014). The detailed docking studies indicate that the identified molecules interact at the reported RNA binding site of N-protein (Kang et al., 2020) and found to have additional interactions that enhance their binding affinity towards the NTD as compared to nucleotides (Fig. 1, S3, and ST 1). The identified molecules targeting the RNA binding pocket were validated using ITC measurements, confirming their high binding affinities towards NTD with low micromolar K_D_ values, ranged from 5.95 - 90 μM (Fig. 2 and Table 1). The NTD-RNA binding fluorescence assay showed high-affinity interaction between NTD and RNA (Luan et al., 2022) (Figure S5). This assay showed that the identified molecules may inhibit the interactions of NTD with the viral genomic RNA as a decrease in fluorescence intensity is observed in presence of small molecules (Fig. 3). The antiviral potential of twelve selected molecules against SARS-CoV-2 was evaluated through cell-based *in vitro* assays after obtaining promising results from docking, biophysical, and fluorescence-based binding inhibition studies. The EC_50_ values of Telmisartan and BMS-189453 were found to be strong, with values ∼1.0 μM and ∼0.98 μM respectively. In contrast, Apatinib and BMS927711 were the least effective (Fig. 4). This study suggests a promising approach for developing antiviral therapy to combat emerging variants of SARS-CoV-2.

Interestingly, most of these small molecules identified in this study are reported as anti-cancerous, anti-diabetic, and some are commercially available as antivirals (Table ST 2). Therefore, this study identified effective anti-SARS-CoV-2 molecules that target the RNA binding pocket of NTD, with EC_50_ values ranging from 0.98-23 μM, that have the potential to be repurposed as monotherapy or combination therapy to fight against SARS-CoV-2 and are expected to be effective against emerging variants. Further studies using relevant animal models and subsequent human clinical trials are required to use any of these molecules as a drug candidate for SARS-CoV-2 treatment.

## Acknowledgements

This work was supported by grant Intensification of Research in High Priority Areas (IRHPA) program of Science and Engineering Research Board (SERB), Department of Science & Technology (DST), Government of India (Grant No.- IPA/2020/000054). Authors are funded by Ministry of Human Resource and Development (MHRD), and Council of Scientific & Industrial Research (CSIR), Government of India.

We thank the participants and contributors to the original research publication. We would like to extend our thanks to the Department of Biosciences and Bioengineering (BSBE), Department of chemistry for providing lab facility, Bioinformatics Centre (BIC), supported by Government of India, (reference number BT/PR40141/BTIS/137/16/2021), the Macromolecular Crystallographic Facility (MCU) for the computer facility, at Indian Institute of Technology Roorkee (IIT Roorkee). We also thank, Indian Veterinary Research Institute, Izatnagar, Bareilly, Uttar Pradesh, India, for BSL-3 facility.

A preprint of this paper is available at bioRxiv; doi: https://doi.org/10.1101/2022.03.12.484092.

## Data Availability Statement

The authors have included the data in the result section of the manuscript and some supporting data are available in supplementary file.

## Supplementary data

Figures S1 to S5 and Tables ST1 and ST2 are available as supplementary data.

