## Supplementary material for "Mechanistic and thermodynamic characterization of antivirals targeting druggable pocket of SARS-CoV-2 nucleocapsid": supplimentry files

### SUPPLEMENTARY DATA

#### *Supplementary results*

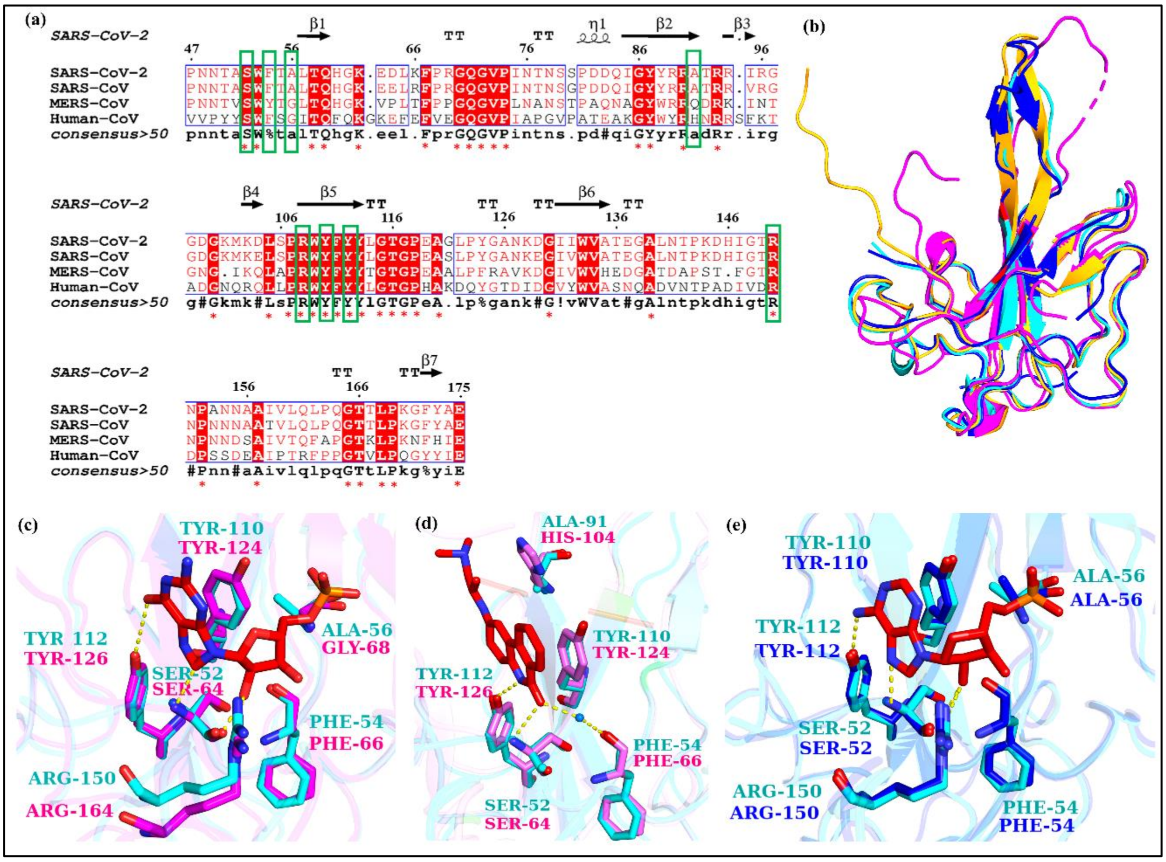


**Supplementary Figure S1. SARS-CoV-2 N-protein is highly conserved at the RNA binding pockets.** (a) Multiple sequence alignment of SARS-CoV-2 N-NTD (PDB ID: 6M3M) with SARS-CoV N-NTD (PDB ID: 2OFZ), MERS-CoV (PDB ID: 4UD1), and HCoV-OC43 (PDB ID 4LI4). Red color stars show conserved residues, and green boxes indicate the residues present in the RNA binding site. (b) The three-dimensional structural alignments of SARS-CoV-2 (Cyan), SARS-CoV (Blue), MERS-CoV (Yellow-orange), and HCoV-OC43 (Magenta). (c) Structural superimposition of GMP binding residues of SARS-CoV-2 (PDB ID: 6M3M) and HCoV-OC43 (PDB ID: 4LM9) with GMP (Red stick). (d) Structural superimposition of GMP binding residues of SARS-CoV-2 (PDB ID: 6M3M) and HCoV-OC43 (PDB ID: 4KJX) with ligand PJ34 (Red stick). (e) Structural superimposition of AMP binding residues of SARS-CoV-2 (PDB ID: 6M3M), SARS-CoV (PDB ID: 2OFZ) with AMP (Red stick), where sticks represent the key RNA binding residues.

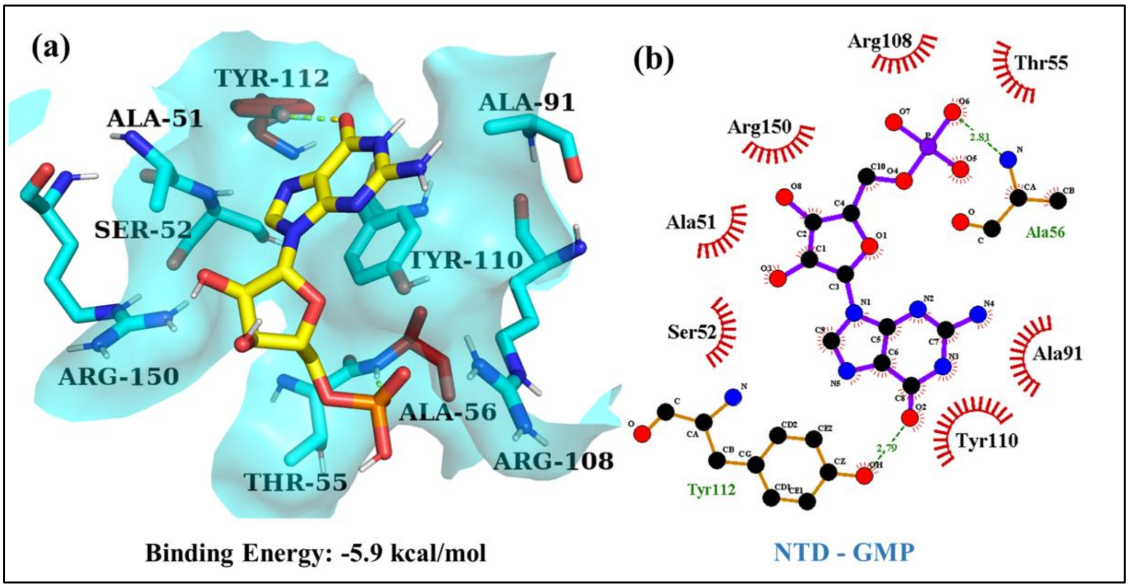


**Supplementary Figure S2.** Detailed schematic representation of NTD-GMP complex. (a) Three-dimensional molecular interaction analysis of GMP with RNA binding pocket of NTD was shown using PyMol. NTD amino acid residues are shown by red and cyan while GMP is in yellow. The surface covered by cyan colour with 60% transparency. Red-coloured residues show H-bond interactions, while cyan-coloured residues show hydrophobic interactions with GMP. The yellow-coloured dotted line guides h-bond interactions. (b) 2D schematic view of molecular interactions of NTD-GMP complex. GMP is represented in purple sticks, while the residues of NTD involved in H-bond interactions are displayed as brown-coloured sticks and green-coloured dotted lines show their H-bond lengths. The red stellate residues represent hydrophobic interactions.

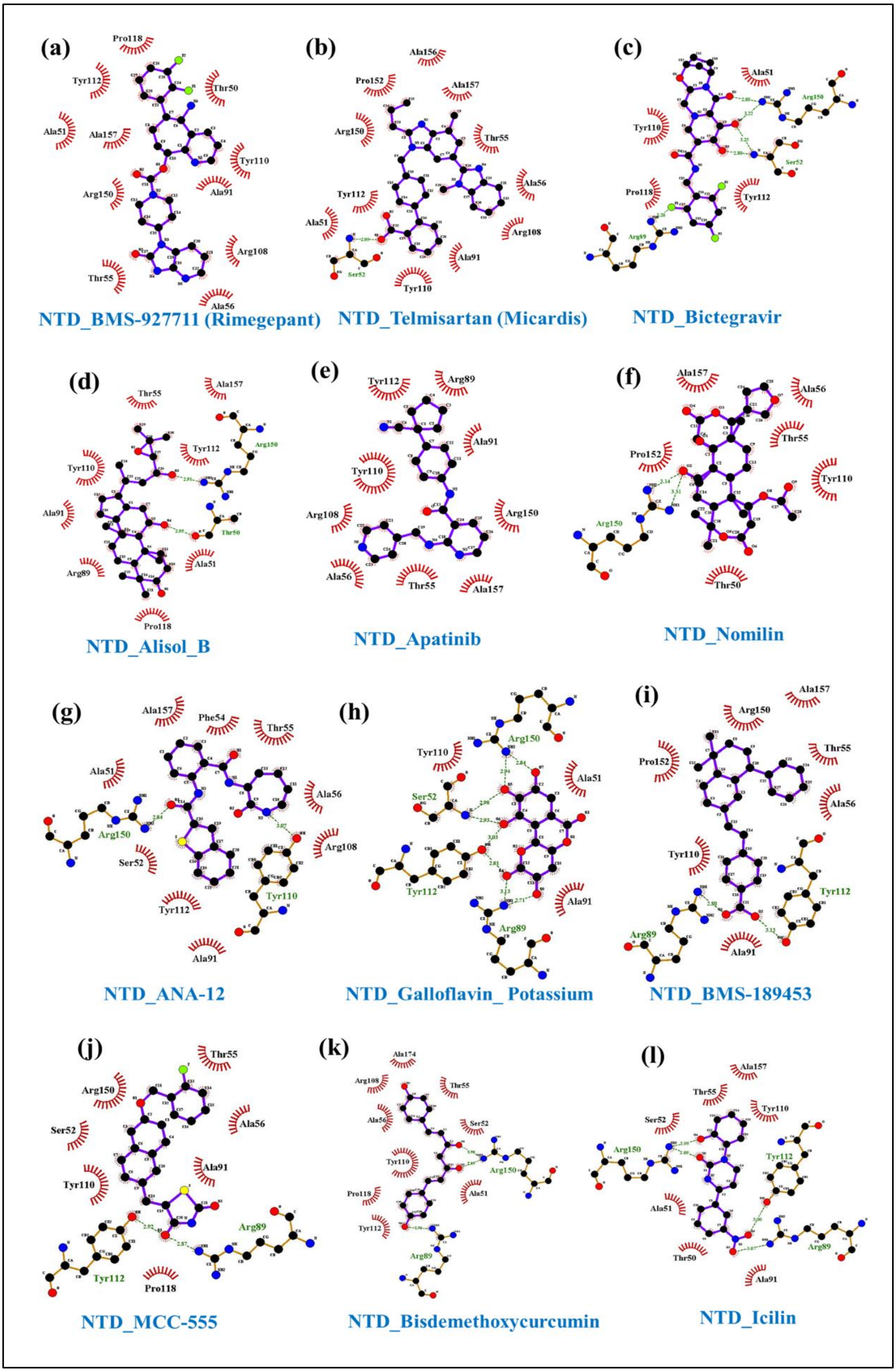
(a)
Pro118
Tyr112
Thr50
Ala51
Ala157
Tyr110
Arg150
Ala91
Arg108
Thr55
Ala56
NTD_BMS-927711 (Rimegepant)
(b)
Ala156
Pro152
Ala157
Arg150
Thr55
Tyr112
Ala56
Ala51
Arg108
Ser52
Ala91
Tyr110
NTD_Telmisartan (Micardis)
(c)
Ala51
Arg150
Tyr110
Ser52
Pro118
Tyr112
Arg89
NTD_Bictegravir
(d)
Ala157
Thr55
Tyr112
Arg150
Tyr110
Thr50
Ala91
Ala51
Arg89
Pro118
NTD_Alisol_B
(e)
Tyr112
Arg89
Ala91
Tyr110
Arg108
Arg150
Ala56
Thr55
Ala157
NTD_Apatinib
(f)
Ala157
Ala56
Thr55
Pro152
Tyr110
Arg150
Thr50
NTD_Nomilin
(g)
Ala157
Phe54
Thr55
Ala51
Ala56
Arg150
Arg108
Ser52
Tyr110
Tyr112
Ala91
NTD_ANA-12
(h)
Tyr110
Arg150
Ser52
Ala51
Tyr112
Ala91
Arg89
NTD_Galloflavin_ Potassium
(i)
Ala157
Arg150
Thr55
Pro152
Ala56
Tyr110
Tyr112
Arg89
Ala91
NTD_BMS-189453
(j)
Thr55
Arg150
Ala56
Ser52
Ala91
Tyr110
Arg89
Tyr112
Pro118
NTD_MCC-555
(k)
Ala174
Arg108
Thr55
Ala56
Ser52
Tyr110
Arg150
Pro118
Ala51
Tyr112
Arg89
NTD_Bisdemethoxycurcumin
(l)
Ala157
Thr55
Tyr110
Ser52
Arg150
Tyr112
Ala51
Arg89
Thr50
Ala91
NTD_Icilin

**Supplementary Figure S3.** Molecular interactions in SARS-CoV-2 NTD-compound complexes in 2D schematic view analysed using LigPlot+. Ligands are represented in purple coloured sticks. The residues of NTD involved in hydrophobic interactions are marked as semi-circled red colour, brown colour sticks represent residues involved in intermolecular H-bond interactions, and green coloured dotted lines show their bond lengths.

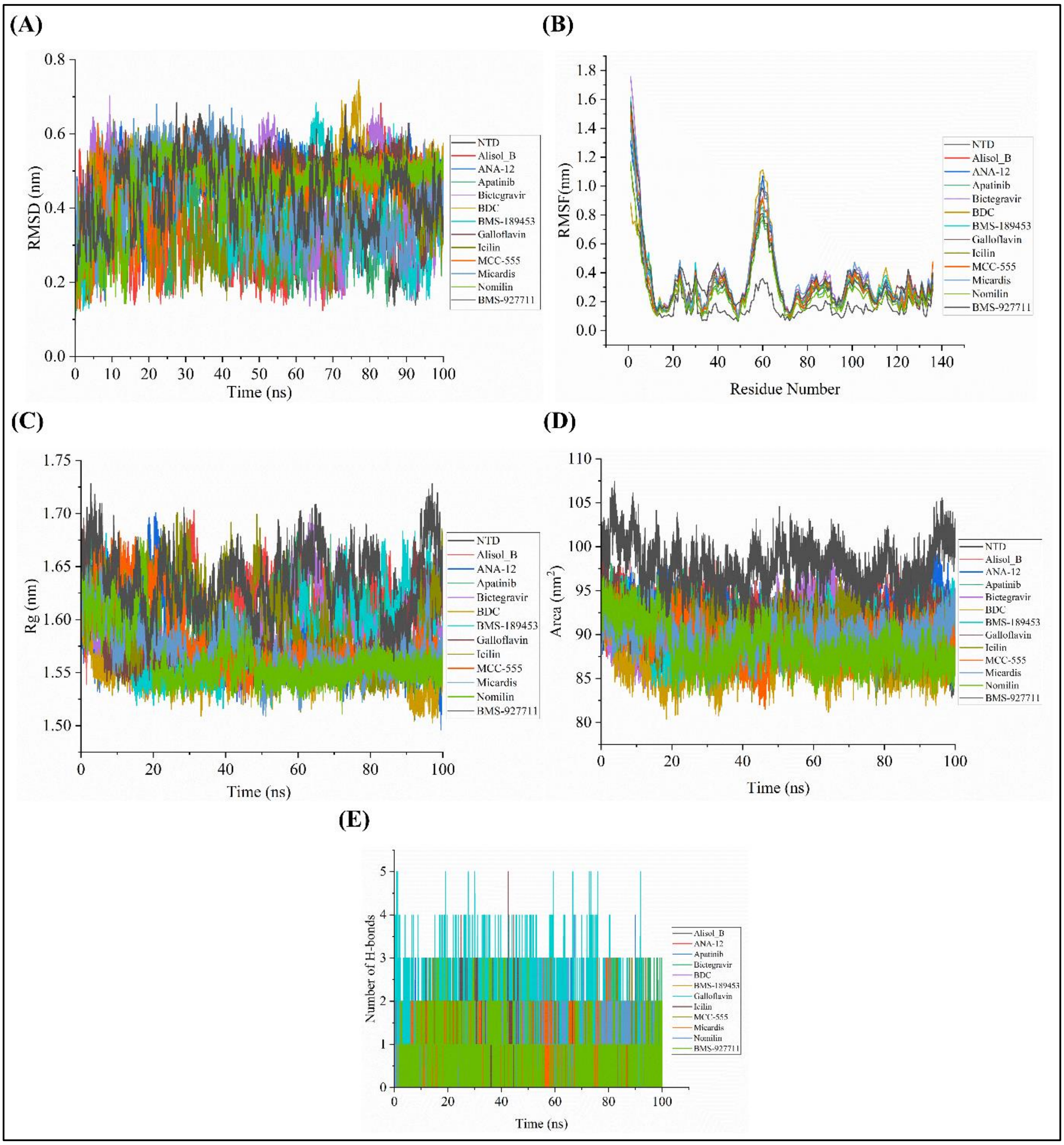


**Supplementary Figure S4**. The schematic representation of the molecular dynamics (MD) simulation studies of NTD in unbound with top twelve selected compounds. (a) RMSD plots (b) RMSF (c) Rg (d) SASA (e) Number of H-bonds formed between NTD and top twelve compounds for 100 ns of MD simulation. The MD simulation trajectories were represented in different colours as NTD (black color), Alisol_B (red), ANA-12 (blue), Apatinib (green), Bictegravir (purple), BDC (mustard), BMS-189453 (cyan), Galloflavin (brown), Icilin (olive), MCC-555 (orange), Micardis (sky blue), Nomilin (grass green), VX-809 (grey).

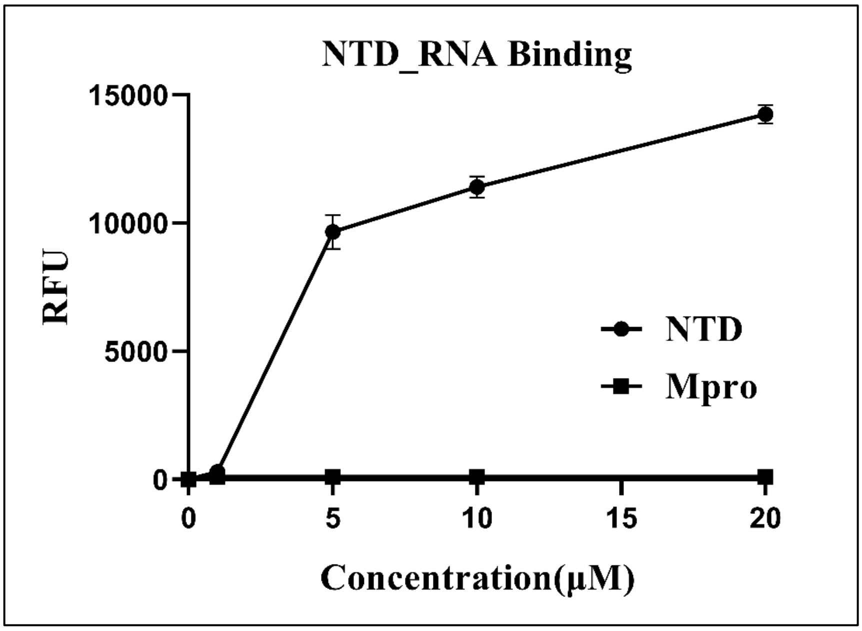


**Supplementary Figure S5.** Fluorescence intensity-based NTD-RNA binding assay. Changes in fluorescence intensity of FAM labelled RNA upon binding to NTD protein. The different concentrations (0µM, 1µM, 5 µM,10 µM, and 20µM) of NTD was titrated with 1nM of FAM-labelled RNA. Error bars represent the standard deviation from duplicate measurements. Different concentrations (0-20µM range) of Mpro (Non- RNA binding protein) taken as negative control.

**Supplementary Table ST1.** Molecular docking of screened compounds against the nucleotide-binding site of NTD protein provides detailed information about binding energy, H-bond & hydrophobic interactions. Here, additional residues are underlined and highlighted.

| Compound | Binding Energy (kcal/mol) | H-Bond Interactions | | Hydrophobic Interactions |
|---|---|---|---|---|
| | | No. of bonds | Residues | |
| FDA Approved compounds | | | | |
| BMS-927711 | -7.9 | 1 | Tyr110 | Thr50, Ala51, Thr55, Ala56, Ala91, Arg108, Tyr112, Pro118, Arg150, Ala157 |
| Telmisartan (MICARDIS) | -7.6 | 2 | Ser52, Tyr112 | Ala51, Thr55, Ala56, Ala91, Arg108, Tyr110, Arg150, Pro152, Ala156, Ala157 |
| Bictegravir | -7.5 | 6 | Ser52, Arg89, | Ala51, Tyr110, Pro118 |

| | | | | |
|---|---|---|---|---|
| | | | Tyr112, Arg150 | |
| Natural compounds | | | | |
| Alisol_B | -7.6 | 2 | Thr50, Arg150 | Ala51, Thr55, Arg89, Ala91, Tyr110, Tyr112, Pro118, Ala157 |
| Apatinib | -7.4 | 1 | Arg89 | Thr55, Ala56, Ala91, Arg108, Tyr110, Tyr112, Arg150, Ala157 |
| Nomilin | -7.1 | 4 | Ala56, Arg150, Ala157 | Thr50, Thr55, Tyr110, Pro152 |
| LOPAC compounds | | | | |
| ANA-12 | -7.4 | 2 | Tyr110, Arg150 | Ala51, Ser52, Phe54, Thr55, Ala56, Ala91, Arg108, Tyr112, Ala157 |
| Galloflavin Potassium | -7.2 | 9 | Ser52, Arg90, Arg89, Tyr112, Arg150 | Ala51, Ala91, Tyr110 |
| BMS-189453 | -6.9 | 2 | Arg89, Tyr112 | Thr55, Ala56, Ala91, Tyr110, Arg150, Asn151, Pro152, Ala157 |
| MCC-555 | -6.9 | 3 | Arg89, Tyr112, Arg150 | Ser52, Thr55, Ala56, Ala91, Tyr110, Pro118 |
| Bisdemethoxycurcumin | -6.8 | 3 | Arg89, Arg150 | Ala51, Ser52, Thr55, Ala56, Arg108, Tyr110, Tyr112, Pro118, Ala174 |
| Icilin | -6.8 | 3 | Arg89, Tyr112, Arg150 | Thr50, Ala51, Ser52, Thr55, Ala91, Tyr110, Ala157 |

**Supplementary Table ST2.** Two-dimensional chemical structure of identified compounds and their reported pharmacological functions

| S. No | Compound Name | Structure | Disease | Function |
|---|---|---|---|---|
| 1. | BMS-927711 (Rimgepant) | 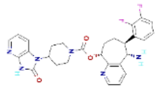 | Preventive treatment of migraine | Antagonist of the CGRP receptor (Jiang et al., 2022), |
| 2. | Telmisartan (MICARDIS) | 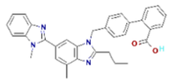 | Treatment of hypertension | Angiotensin II type 1 (AT1) receptor antagonist used in the management of hypertension (Ruilope and Schumacher, 2012), |
| 3. | Bictegravir | 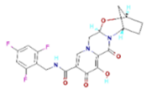 | Treatment of HIV infection | Antiviral drug (Gaur et al., 2021; |

| | | | | |
|---|---|---|---|---|
| | | | | Wohl et al., 2019) |
| 4. | Alisol_B | 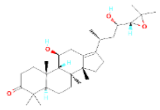 | Treatment of cancer (Breast cancer) | Anticancerous (Zhang et al., 2017) |
| 5. | Apatinib | 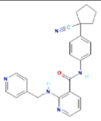 | Treatment of cancer (Breast cancer, lung cancer) | Antineoplastic Agents, Protein Kinase Inhibitors (Sun et al., 2020) |
| 6. | Nomilin | 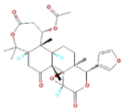 | Inhibition of tumor progression by naturally occurring terpenoids | Triterpenoid present in common edible citrus fruits with putative anticancer properties (Giofrè et al., 2021) |

| | | | | |
|---|---|---|---|---|
| 7. | ANA-12 | 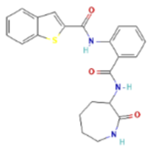 | Anti-depressant | Dopamine D3 receptor (Ribeiro et al., 2020)’ |
| 8. | Galloflavin_Potassium | 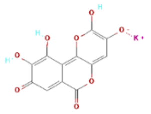 | HIV-1 integrase inhibitors | Antiviral against HIV-1 and a Lactate dehydrogenase inhibitor (Hong et al., 1998; Manerba et al., 2012) |
| 9. | BMS-189453 | 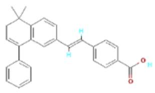 | Treatment of arthritis | Synthetic retinoid receptor antagonist (Amati et al., 2010) |
| 10. | MCC-555 | 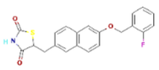 | Anti-proliferative, Anti-diabetic | Peroxisome proliferator-activated |

| | | | | |
|---|---|---|---|---|
| | | | | receptor ligand (Sun et al., 2009) |
| 11. | Bisdemethoxycurcumin | 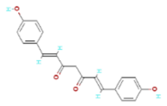 | Anti-inflammatory, Anti-diabetic | Antioxidant (Kou et al., 2013; Ponnusamy et al., 2012) |
| 12. | Icilin | 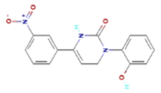 | Cooling agent, Treatment of diseases digestive system | Calcium Channel Agonists (Selescu et al., 2013) |
